# The ALT pathway generates telomere fusions that can be detected in the blood of cancer patients

**DOI:** 10.1101/2022.01.25.477771

**Authors:** Francesc Muyas, Manuel José Gómez Rodriguez, Isidro Cortes-Ciriano, Ignacio Flores

## Abstract

Telomere fusions (TFs) can trigger the accumulation of diverse genomic rearrangements and the acquisition of oncogenic alterations leading to malignant transformation and resistance to chemotherapy. Despite their relevance in tumour evolution, our understanding of the patterns and consequences of TFs in human cancer remains limited. Here, we have characterized the rates and spectrum of somatic TFs across >30 cancer types using whole-genome sequencing data. TFs are pervasive in human tumours with rates varying markedly across and within cancer types. In addition to end-to-end fusions, we find novel patterns of TFs that we mechanistically link to the activity of the alternative lengthening of telomeres (ALT) pathway. We show that TFs can be detected in the blood of cancer patients, which enables cancer detection with high specificity and sensitivity even for early-stage tumours and cancer types for which early detection remains a high unmet clinical need, such as pancreatic cancer and brain tumours. Overall, we report a novel genomic footprint that enables characterization of the telomere maintenance mechanism of tumours and liquid biopsy analysis, which has implications for early detection, prognosis, and treatment selection.

## Introduction

Telomeres are nucleoprotein complexes composed of telomeric TTAGGG repeats and telomere binding proteins that prevent the recognition of chromosome ends as sites of DNA damage^1^. The replicative potential of somatic cells is limited by the length of telomeres, which shorten at every cell division due to end-replication losses. Most human cancers maintain the length of telomeres above a critical threshold by re-expressing telomerase through diverse mechanisms^2^, including activating *TERT* promoter mutations^3,4^ and enhancer hijacking^5^. In other cancers, in particular those of mesenchymal and neuroendocrine origin, telomeres are elongated by the alternative lengthening of telomeres (ALT) pathway, which relies on recombination^6^. Telomere attrition can result in senescence or the ligation of chromosome ends to form dicentric chromosomes, which are observed as chromatin bridges during anaphase^7^. The resolution of chromosome bridges caused by telomere fusions (TFs) can increase genomic instability and the acquisition of oncogenic alterations involved in malignant transformation and resistance to chemotherapy through diverse mechanisms, including chromothripsis and breakage-fusion-bridge cycles^8–12^.

Despite their importance in tumour evolution, the patterns and consequences of TFs remain largely uncharacterized, in part due to technical challenges. TFs have been traditionally detected by inspection of chromosome bridges in metaphase spreads, an approach that is not scalable and has low resolution^13–15^. In recent years, the study of TFs has relied on PCR-based methods using primers annealing to subtelomeric regions^16,17^, which are limited to detect TFs distantly located from subtelomeres since PCR efficiency decreases as the amplicon size increases^18^. To overcome these limitations, we have developed computational methods to detect TFs using whole-genome sequencing (WGS) data. Here, we report the rates and patterns of TFs across >2,000 tumours spanning >30 cancer types. In addition, we report the discovery of a novel type of TF which we mechanistically link with the ALT pathway. We also demonstrate that ALT and non-ALT tumours generate TFs that are detected in the blood of cancer patients with high sensitivity and specificity for liquid biopsy analysis.

## Results

### Pan-cancer landscape of telomere fusions

To detect TFs in sequencing data, we developed TelFusDetector (Fig. 1a). In brief, TelFusDetector identifies sequencing reads containing at least two consecutive TTAGGG and two consecutive CCCTAA telomere sequences, allowing for mismatches to account for the variation observed in telomeric repeats in humans^19,20^ (Fig. 1 and Supplementary Fig. 1; see Methods for a detailed description of the algorithm, including QC steps). We first scanned the human reference genome to identify regions containing telomere fusion-like patterns, which could be misinterpreted as somatic (Methods). Our analysis revealed the relic of an ancestral fusion in chromosome 2^21^, and a region in chromosome 9 containing 2 sets of telomeric repeats flanked by high complexity sequences, which we term “chromosome 9 endogenous fusion” (Supplementary Table 1).

**Figure 1.**
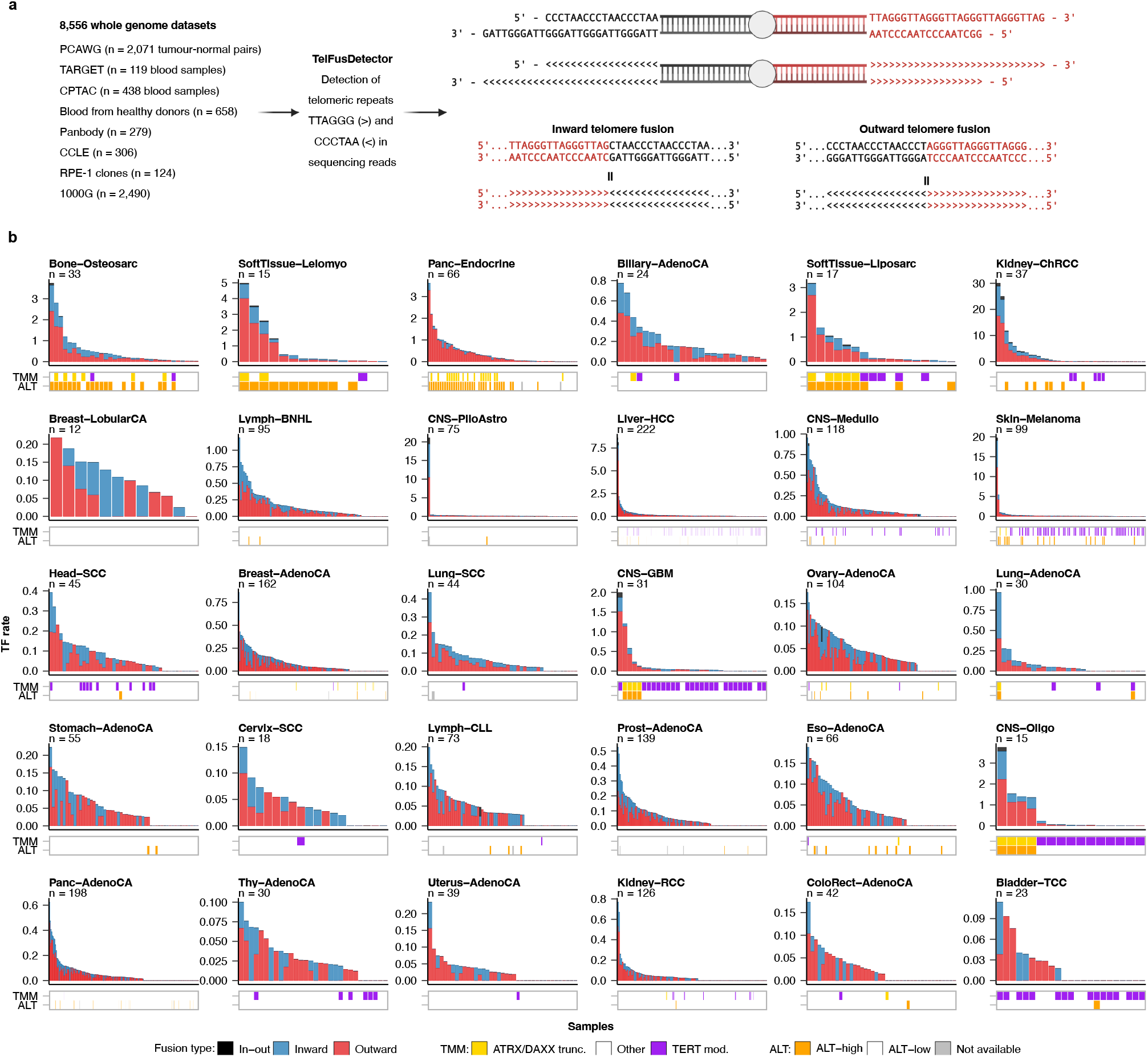
Landscape of telomere fusions in cancer. **a**, Overview of the study design and schematic representation of the two types of telomere fusions (TFs) identified by TelFusDetector. **b**, TF rates across 30 cancer types from PCAWG. Outward fusions are shown in red and inward in blue. Read pairs where a read in the pair contained an inward TF and the other an outward TF, which would be consistent with a circular DNA element, are indicated as in-out TFs and shown in grey. The colour of the boxes below the bar plots represent the telomere maintenance mechanism (TMM) predictions reported by Nonneville et al.^22^ and Sieverling et al.^19^. Only cancer types with at least 10 tumours are shown. ATRX/DAXXtrunc, tumours with inactivating point mutations, frameshift indels and structural variants in ATRX or DAXX; TERT mod., tumours with activating mutations in *TERT*. The abbreviations used for the cancer types are as follows: Biliary-AdenoCA, biliary adenocarcinoma; Bladder-TCC, bladder transitional cell carcinoma; Bone-Benign, bone cartilaginous neoplasm, osteoblastoma and bone osteofibrous dysplasia; Bone-Epith, bone neoplasm, epithelioid; Bone-Osteosarc, sarcoma, bone; Breast-AdenoCA, breast adenocarcinoma; Breast-DCIS, breast ductal carcinoma in situ; Breast-LobularCA, breast lobular carcinoma; Cervix-AdenoCA, cervix adenocarcinoma; Cervix-SCC, cervix squamous cell carcinoma; CNS-GBM, central nervous system glioblastoma; CNS-Oligo, CNS oligodenroglioma; CNS-Medullo, CNS medulloblastoma; CNS-PiloAstro, CNS pilocytic astrocytoma; ColoRect-AdenoCA, colorectal adenocarcinoma; Eso-AdenoCA, esophagus adenocarcinoma; Head-SCC, head-and-neck squamous cell carcinoma; Kidney-ChRCC, kidney chromophobe renal cell carcinoma; Kidney-RCC, kidney renal cell carcinoma; Liver-HCC, liver hepatocellular carcinoma; Lung-AdenoCA, lung adenocarcinoma; Lung-SCC, lung squamous cell carcinoma; Lymph-CLL, lymphoid chronic lymphocytic leukemia; Lymph-BNHL, lymphoid mature B-cell lymphoma; Lymph-NOS, lymphoid not otherwise specified; Myeloid-AML, myeloid acute myeloid leukemia; Myeloid-MDS, myeloid myelodysplastic syndrome; Myeloid-MPN, myeloid myeloproliferative neoplasm; Ovary-AdenoCA, ovary adenocarcinoma; Panc-AdenoCA, pancreatic adenocarcinoma; Panc-Endocrine, pancreatic neuroendocrine tumor; Prost-AdenoCA, prostate adenocarcinoma; Skin-Melanoma, skin melanoma; SoftTissue-Leiomyo, leiomyosarcoma, soft tissue; SoftTissue-Liposarc, liposarcoma, soft tissue; Stomach-AdenoCA, stomach adenocarcinoma; Thy-AdenoCA, thyroid low-grade adenocarcinoma; and Uterus-AdenoCA, uterus adenocarcinoma.

To characterize the patterns and rates of somatic TFs across diverse cancer types, we applied TelFusDetector to 2071 matched tumour and normal sample pairs from the Pan-Cancer Analysis of Whole Genomes (PCAWG) project that passed our QC criteria (Methods and Supplementary Fig. 1). To enable comparison of the relative number of TFs across samples, we computed a TF rate for each tumour after correcting for tumour purity, sequencing depth, and read length (Methods, Supplementary Fig. 2 and Supplementary Table 2). We identified two distinct TF patterns, which differ in the relative position of the sets of TTAGGG and CCCTAA repeats (Fig. 1a). A first pattern, which we term “inward TF”, is characterized by 5’-TTAGGG-3’ repeats followed by 5’-CCCTAA-3’ repeats, which is the expected genomic footprint of end-to-end TFs^1,13^. Unexpectedly, we also found a second pattern characterized by 5’-CCCTAA-3’ repeats followed by 5’-TTAGGG-3’ repeats, which we term “outward TF” (Fig. 1a), and, to the best of our knowledge, represents a novel class of structural variation. In addition, we found read pairs where a read in the pair contained an inward TF and the other an outward TF, which would be consistent with a circular DNA element, and thus we term these circular (in-out) TFs.

Both outward and inward TFs were detected across diverse cancer types, but rates varied markedly within and across tumour types (Fig. 1b). The highest TF rates were observed in osteosarcomas (Bone-Osteosarc), leiomyosarcomas (SoftTissue-Leiomyo), and pancreatic neuroendocrine tumours (Panc-Endocrine). The lowest frequencies were observed in thyroid adenocarcinomas (Thy-AdenoCA), renal cell-carcinomas (Kidney-RCC), and uterine adenocarcinomas (Uterus-AdenoCA). These results indicate that somatic TFs, including the novel type of outward fusions we report here, are pervasive across diverse cancer types.

### The ALT pathway is mechanistically linked with the formation of telomere fusions

Next, we sought to determine the molecular mechanisms implicated in the generation of TFs. To this aim, we regressed the observed rates of TFs on the mutation status of *ATRX* and *DAXX*, telomere content, point mutations and structural variants in the *TERT* promoter, expression values of *TERT* and TERRA, and a binary category indicating the ALT status of each tumour predicted using two previously published classifiers^19,22^ (Methods). Our analysis revealed a strong association between the activation of the ALT pathway and the rate of TFs, with the strongest effect size observed for outward TFs (*P* < 0.05; Fig. 2a,b and Supplementary Fig. 3a). However, alterations of the *TERT* promoter were negatively correlated with both inward and outward fusion rates, with the highest effect size obtained for outward TFs (Fig. 2a). The association of TF rates with telomere content and TERRA expression was also significant, although of a modest effect size (*P* < 0.001, ANOVA; Supplementary Table 3).

**Figure 2.**
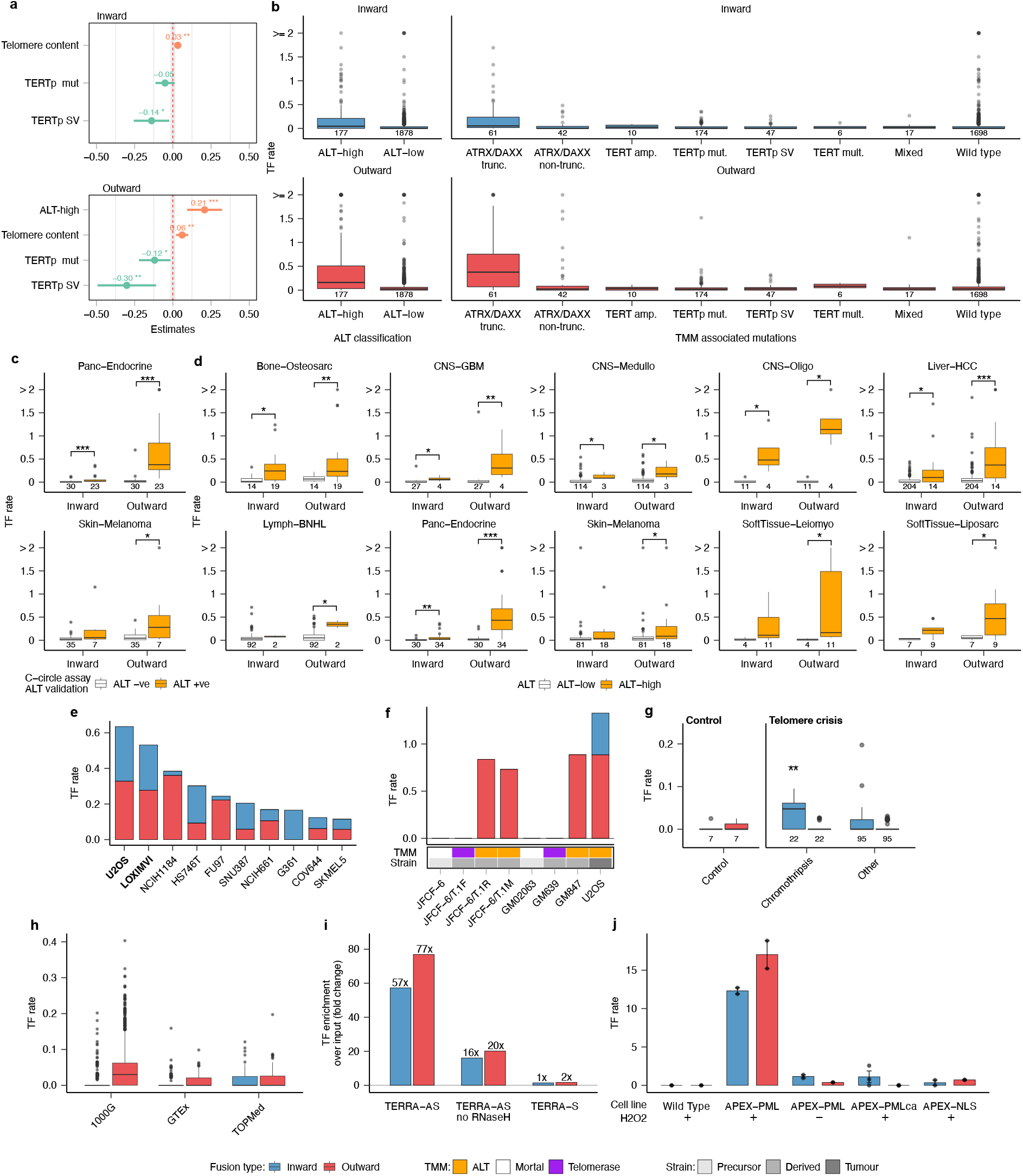
TFs are generated by the activity of the ALT pathway. **a**, Coefficient values estimated using linear regression analysis and variable selection for the covariates with the strongest positive and negative association with TF rates. For this analysis we used the ALT status classification reported by de Nonneville et al.^22^. **b**, Rates of inward and outward fusions in PCAWG tumours grouped by ALT status predictions (de Nonneville et al. ^22^), and TMM-associated mutations (Sieverling et al.^19^). **c**, Comparison of TF rates between tumours positive and negative for the C-circle assay. **d**, TF rates estimated for PCAWG tumours grouped by ALT status predictions across selected cancer types. Only cancer types with significant differences in the number of TFs based on ALT status are shown. **e**, Top 10 cancer cell lines from the CCLE with the highest TF rates. ALT cell lines are indicated in bold type. **f**, TF rates in mortal cell strains before and after transformation by mechanisms requiring telomerase or ALT. **g**, TF rates detected in RPE-1 cell lines before (control) and after induction of telomere crisis with doxycycline. **h**, TF rates detected in 1000G, GTEx and TOPMed samples. **I**, Fold changes in TF rates over the control estimated using CHIRT-seq data from mouse embryonic stem cells treated with TERRA-AS, TERRA-AS without Rnase H, and TERRA-S. **j**, TF rates estimated using AlaP data generated using different conditions of APEX knock-in and peroxidase (H_2_O_2_). In all panels \*\*\**P* < 0.001; \*\**P* < 0.01; \**P* < 0.05, Wilcoxon rank-sum test after FDR correction. Box plots show the median, first and third quartiles (boxes), and the whiskers encompass observations within a distance of 1.5x the interquartile range from the first and third quartiles. ALT −ve, ALT negative; ALT +ve, ALT positive; ATRX/DAXXtrunc, tumours with inactivating point mutations, frameshift indels and structural variants in ATRX or DAXX; ATRX/DAXX non-trunc, tumours without inactivating mutations in ATRX or DAXX; TERT amp., tumours with at least 6 additional *TERT* copies over the tumour ploidy; TERTp mut., tumours with activating mutations in the *TERT* promoter region; TERTp SV, with a structural variation (SV) breakpoint on the plus strand 20 kb upstream of *TERT*; TERT mult., tumours with multiple *TERT* alterations. Mixed, tumours with both an activation *TERT* alteration and an ATRX/DAXX alteration.

To investigate the association between the ALT pathway and TF formation, we first compared the rate of TFs between tumours positive and negative for C-circles, an ALT marker^19,22^. For this analysis, we focused on published data from 42 skin melanomas and 53 pancreatic neuroendocrine tumours that are also part of the PCAWG cohort. ALT tumours showed significantly higher rates of TFs in the pancreatic neuroendocrine tumour set (*P* < 0.001, two-tailed Mann-Whitney test; Fig. 2c). A similar trend was observed for skin melanomas, although only outward fusion rates reached significance (Fig. 2c). Next, we extended this analysis to the entire cohort by comparing the TF rates between tumours with high and low ALT-probability scores (ALT-low vs ALT-high)^22^ on a per cancer type basis (Fig. 2d). Overall, TF rates, in particular outward TFs, were significantly higher in cancer types classified as ALT-high (FDR-corrected *P* < 0.1, two-tailed Mann-Whitney test; Fig. 2d and Fig 1b).

To test whether TF fusions are enriched in ALT cancers, we analysed whole-genome sequencing data from 306 cancer cell lines from the Cancer Cell Line Encyclopedia (CCLE)^23^. Consistent with our observations in primary tumours, cell lines used as models of ALT, such as the osteosarcoma cell line U2OS and the melanoma cell line LOXIMVI, showed the highest rates of both inward and outward TFs (Fig. 2e and Supplementary Fig. 1i). Analysis of PacBio long-read sequencing data from the ALT breast cancer cell line SK-BR-3^24^ also revealed an enrichment of outward TFs in this line as compared to the non-ALT cell lines COLO289T, HCT116, KM12, SW620 and SW837, for which long-read sequencing data were also available^25,26^ (Supplementary Fig. 3b-c; see Supplementary Fig. 4 for examples).

To assess whether TFs are specifically associated with the ALT pathway, we analysed the genomes of mortal cell strains before and after transformation by mechanisms requiring telomerase or ALT^27^. The genomes of parental mortal strains JFCF-6 and GM02063, as well as telomerase-positive strains JFCF-6/T.1F and GM639, did not contain outward TFs (Fig. 2f). In contrast, ALT-derived strains JFCF-6/T.1R, JFCF-6/T.1M and GM847 showed a comparable outward TF fusion rate to the prototypical ALT cell line U2OS (Fig. 2f). Therefore, the presence of outward TFs in ALT derived-strains but not in telomerase-positive strains indicates that ALT activation leads to the formation of outward TFs. In addition, we analysed TF rates in whole-genome sequencing data from hTERT-expressing retinal pigment epithelial (RPE-1) cells sequenced after the induction of telomere crisis using a dox-inducible dominant negative allele of *TRF2*^9,10^. Compared to the control samples sequenced before induction of telomere crisis, we detected a high rate of inward TFs, which is consistent with the presence of end-to-end fusions (*P* < 0.05, two-tailed Mann-Whitney test; Fig. 2g and Supplementary Fig. 1j), thus lending further support to the mechanistic association between ALT activity and the formation of outward TFs. Consistent with the activation of the ALT pathway in cells immortalized by the Epstein-Barr virus *in vitro*^28,29^, we also detected high rates of outward TFs in 2490 Epstein–Barr virus-immortalized B cell lines from the 1000G project (Fig. 2h and Supplementary Fig. 1).

To further test the association between TFs and ALT activity, we used Random Forest classification to predict the ALT status of tumours using the rates and features of TFs as covariates, and the set of tumours with C-circle assay data as the training set (Methods). Variable importance analysis using the best performing classifier (AUC=0.93) identified variables encoding the rate and breakpoint sequences of TFs as the most predictive features, followed by the fraction of the telomere variant repeats (TVR) GTAGGG and CCCTAG, which are enriched in ALT tumours^30^, in sequencing reads with TFs (Supplementary Fig. 5 and Methods).

Together, these results mechanistically link the activity of the ALT pathway with the generation of somatic TFs. Therefore, we term inward and outward fusions ALT-associated TFs (ALT-TFs).

### ALT-TFs bind to TERRA and localize to APBs

We next sought to determine the association of ALT-TFs with molecules involved in telomere maintenance and their cellular localization. Our regression expression analysis of the PCAWG data set indicates that tumours enriched in ALT-TFs present elevated levels of TERRA, a long non-coding RNA transcribed from telomeres^31,32^. Previous genomic and cytological studies demonstrated a preferential association of TERRA transcripts to telomeres^33^. To assess whether TERRA also associates with ALT-TFs, we searched for inward and outward ALT-TFs in reads containing TERRA-binding sites. Specifically, we analysed reads from CHIRT-seq, an immunoprecipitation protocol that specifically captures TERRA-binding sites using an anti-sense biotinylated TERRA transcript (TERRA-AS) as bait^34^. Targets of the TERRA-AS bait are then treated with RNase H to elute DNA containing TERRA binding sites followed by sequencing. By analysing CHIRT-seq data sets from mouse embryonic stem cells (mESCs)^34^, we observed a 57-fold and 77-fold enrichment of inward and outward ALT-TFs, respectively, over the input using the TERRA-AS oligo probe (Fig. 2i). However, a modest enrichment was observed when the TERRA-AS not treated with RNase H or the TERRA sense transcript (TERRA-S) were used (Fig. 2i). These results thus indicate that TERRA binds to inward and outward ALT-TFs.

TERRA transcripts can be found in a subtype of promyelocytic leukaemia nuclear bodies (PML-NB) termed ALT-associated PML-Bodies (APBs)^35^. Because ALT-TFs bind to TERRA, we hypothesized that inward and/or outward ALT-TFs might locate to APBs. Given that PML-NBs, including APBs, are insoluble^36^, a standard ChIP-seq protocol for PML cannot be used to analyse whether ALT-TFs are present in APBs. To overcome PML-NBs accessibility problems, Kurihara *et al*.^37^ recently developed an assay called ALaP, for APEX-mediated chromatin labelling and purification by knocking in APEX, an engineered peroxidase, into the *Pml* locus to tag PML-NB partners in an H_2_O_2_-dependent manner. Applying ALaP in mESCs, PML-NBs bodies were found to be highly enriched in ALT-related proteins, such as DAXX and ATRX, as well as in telomere sequences. Here, to test our hypothesis, we searched for ALT-TFs in ALaP genomic pull-downs and found a strong enrichment of both inward and outward ALT-TFs (*P* < 0.05, two-tailed Mann-Whitney test; Fig. 2j). In addition, we found that negative controls, *i.e*., APEX-PMLs not-treated with H_2_O_2_ or APEX variants that do not form PML-NBs, rarely contain ALT-TFs (Fig. 2j). Therefore, these results indicate that APBs are a preferential location for ALT-TFs.

### Short DNA fragments contain ALT-TFs

In addition to APBs, another feature of ALT positive cells is their elevated levels of extrachromosomal telomeric DNA (ECT-DNA). Interestingly, most ECT-DNAs in ALT positive cells localize to APBs^38^. As ALT-TFs also localize to APBs, it is conceivable that ECT-DNAs exert as substrates for the formation of ALT-TFs. If this was the case, the formation of ALT-TFs would result from the fusion of short ECT-DNA fragments rather than chromosomes. To test this hypothesis, we inferred the fragment size for read pairs with ALT-TF or chromosome 9 endogenous fusions in which both mates support the same breakpoint sequence (Methods). We found a significant enrichment of ALT-TFs in DNA fragments shorter than the average library insert size in a set of cancer types with high ALT-TF rates, such as melanomas, osteosarcomas, and glioblastomas (FDR-corrected *P* < 0.1; Chisquare test; Supplementary Table 4). Complementary analyses performed on mESC PML-NB sequencing data^37^ indicated that 26% of ALT-TF-containing reads were significantly shorter than the average read length, while this happened for around 4% of the reads mapping outside telomeres. In addition, ALT-TF-containing fragments were three times shorter than the average fragment size in the library (Supplementary Fig. 6 and Methods). Together, these results indicate that ALT-TFs might originate from the fusion of small DNA fragments.

### Sequence specificity at the telomere fusion point

We next analysed the set of sequences at the fusion point in ALT-TFs detected in PCAWG tumours. ALT-TFs with breakpoint sequences in the set of all possible circular permutations of TTAGGG and CCCTAA sequences were classified as pure (59% of ALT-TFs), whereas ALT-TFs with complex breakpoint sequences longer than 12bp were classified as alternative (41%; Fig. 3a, Supplementary Table 5 and Methods). In pure ALT-TFs, we detected the entire set of possible permutations of telomere repeat motifs at fusion breakpoints, but not at similar frequencies (*P* < 0.05; chi-square test; Fig. 3b). In the case of outward ALT-TFs, the breakpoint sequence ^5’^…CCCTAACCC**TA**GGGTTAGGG…^3’^ was the most abundant (22% of pure ALT-TFs) followed by ^5’^…CCCTAACCC**TTA**GGGTTAGGG…^3’^ (16%). Interestingly, these two breakpoint sequences can be generated by the ligation of 7 and 4 combinations of telomeric repeats, respectively, while the other breakpoint sequences detected can only be generated by the combination of two specific telomeric repeat sequences (Fig. 3b and Supplementary Fig. 7). In addition, these two sequences are the only ones in the entire set of breakpoint sequences in outward ALT-TFs with microhomology at the fusion point. In inward ALT-TFs, the sequences ^5’^…TTAGGG**TTAA**CCCTAA…^3’^ and ^5’^…TTAGGG**TAA**CCCTAA…^3’^ were the most abundant (14% each). We also detected the TTAGCTAA sequence in 7% of pure ALT-TFs, which could be generated by the end-to-end fusion of two telomeres considering that … TTAG^3’^ is the most common terminal sequence at human telomeres^1^. Similar to outward ALT-TFs, TTAA and TAA are the only breakpoint sequences in inward fusions that can be created by the ligation of several combinations of telomeric repeats and contain microhomology at the fusion junction (Fig. 3b). As microhomology facilitates ligation, we conclude that microhomology at the fusion point also contributes to explain differences in the frequency of specific breakpoint sequences in both outward and inward ALT-TFs.

**Figure 3.**
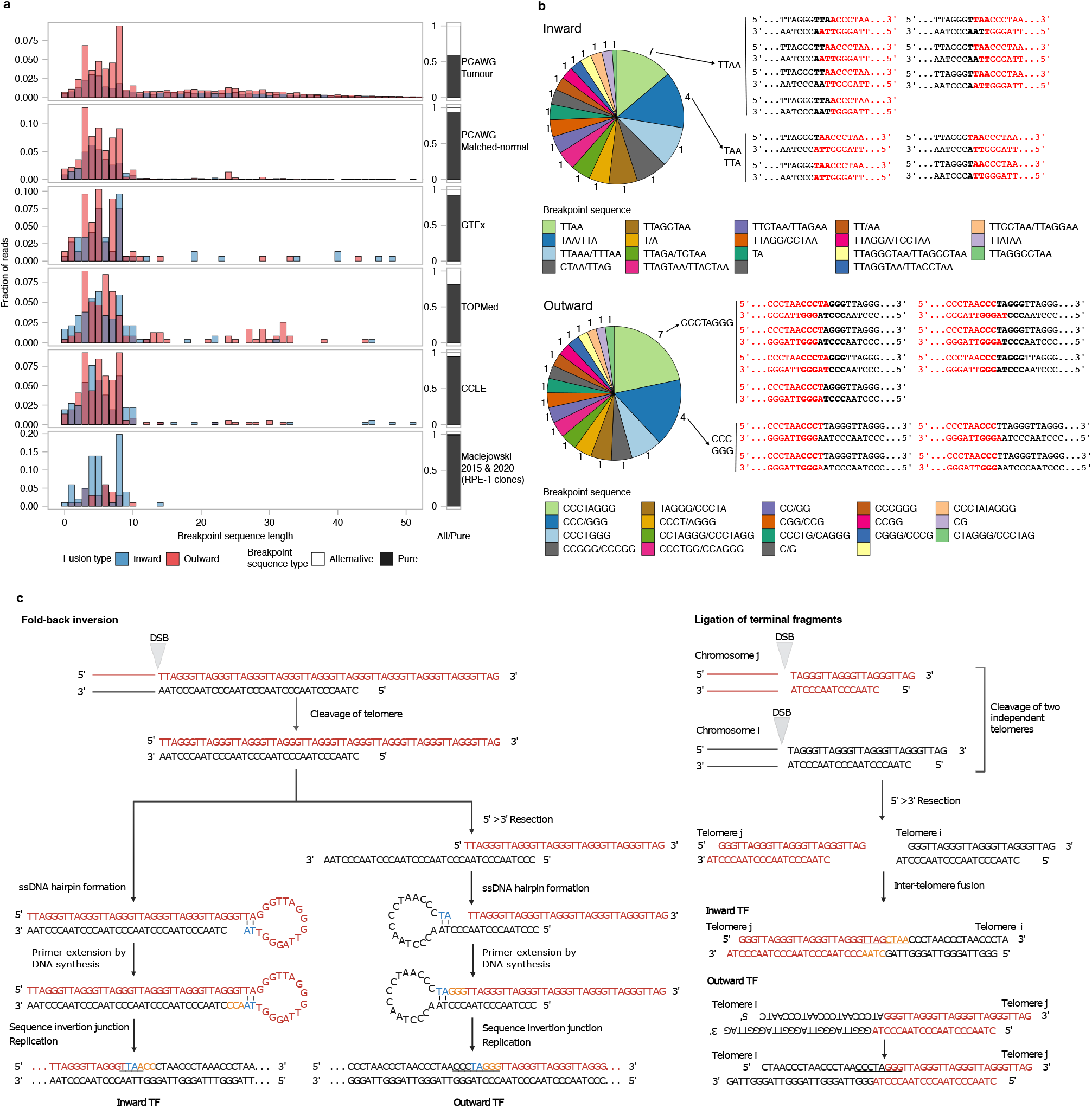
Mechanisms of ALT-TF formation. **a**, Breakpoint sequence length distribution. Inward and outward fusions are shown in blue and red, respectively. The bars on the right show the fraction of ALT-TFs classified as pure (black) or alternative (white). **b**, Pie chart showing the distribution of the distinct breakpoint sequences observed in pure ALT-TFs in PCAWG tumours. The numbers around the pie charts represent the number of combinations of circular permutations of telomeric repeat motifs that can generate each breakpoint sequence. The legend reports the breakpoint sequences in both strands unless they are identical (e.g., TTAA). The most represented breakpoint sequences are indicated. **c**, Proposed mechanisms for ALT-TF formation. ALT-TFs are generated through an intra-telomeric fold-back inversion after a doublestrand break (left), or by the ligation of terminal telomere fragments after double-strand breaks (right).

### ALT-TFs are generated through the repair of double-strand breaks by an intra- or an inter-telomeric mechanism

Our previous analysis suggests that ALT-TFs are generated at APBs preferentially when telomeric fragments with microhomology in their ends fuse. Therefore, we postulate two non-exclusive mechanisms of ALT-TF formation (Fig. 3c and Supplementary Fig. 8). First, a double-strand break in a telomere can be repaired through an intra-telomeric fold-back inversion. Specifically, end resection of a double-strand break would facilitate the formation of a hairpin loop when the 3’ end of a telomere strand folds back to anneal its complementary strand through microhomology. Then, DNA synthesis would fill the gap to complete the capping of the hairpin. Finally, replication of the hairpin would create an inward or an outward ALT-TF depending on the 3’ end telomeric strand that folds back: …(TTAGGG)_n_…^3’^ fold-back would create an inward ALT-TF and …(CCCTAA)_n_…^3’^ fold-back would create an outward ALT-TF. Secondly, ALT-TFs can also be generated through the ligation of the terminal fragments upon double-strand DNA breaks in telomeres (Fig. 3c and Supplementary Fig. 8). Specifically, an inter-telomeric mechanism would occur when two telomeres covalently fuse in ^5’^ …(TTAGGG)_n_…-…(CCCTAA)_n_…^3’^ orientation to create an inward ALT-TF, or in ^5’^ …(CCCTAA)_n_…-…(TTAGGG)_n_…^3’^ orientation to create an outward ALT-TF. Outward ALT-TFs are only feasible when telomeric fragments join from the broken ends produced after telomere trimming (Fig. 3c).

### ALT-TFs are detected in blood and enable cancer detection

Given the high rate of ALT-TFs observed in tumours of diverse origin, we hypothesized that ALT-TFs could also be detected in blood samples and used as biomarkers for liquid biopsy analysis. To test this hypothesis, we applied TelFusDetector to blood samples from PCAWG (n=1604), the Genotype-Tissue Expression (GTEx; n=255) project and the Trans-Omics for Precision Medicine program (TOPMed; n=304) (Methods). Overall, blood samples from cancer patients showed a significantly higher rate of ALT-TFs, especially of the outward type (FDR-corrected *P* < 0.1, twotailed Mann-Whitney test; Fig. 4a and Supplementary Fig. 9).

**Figure 4.**
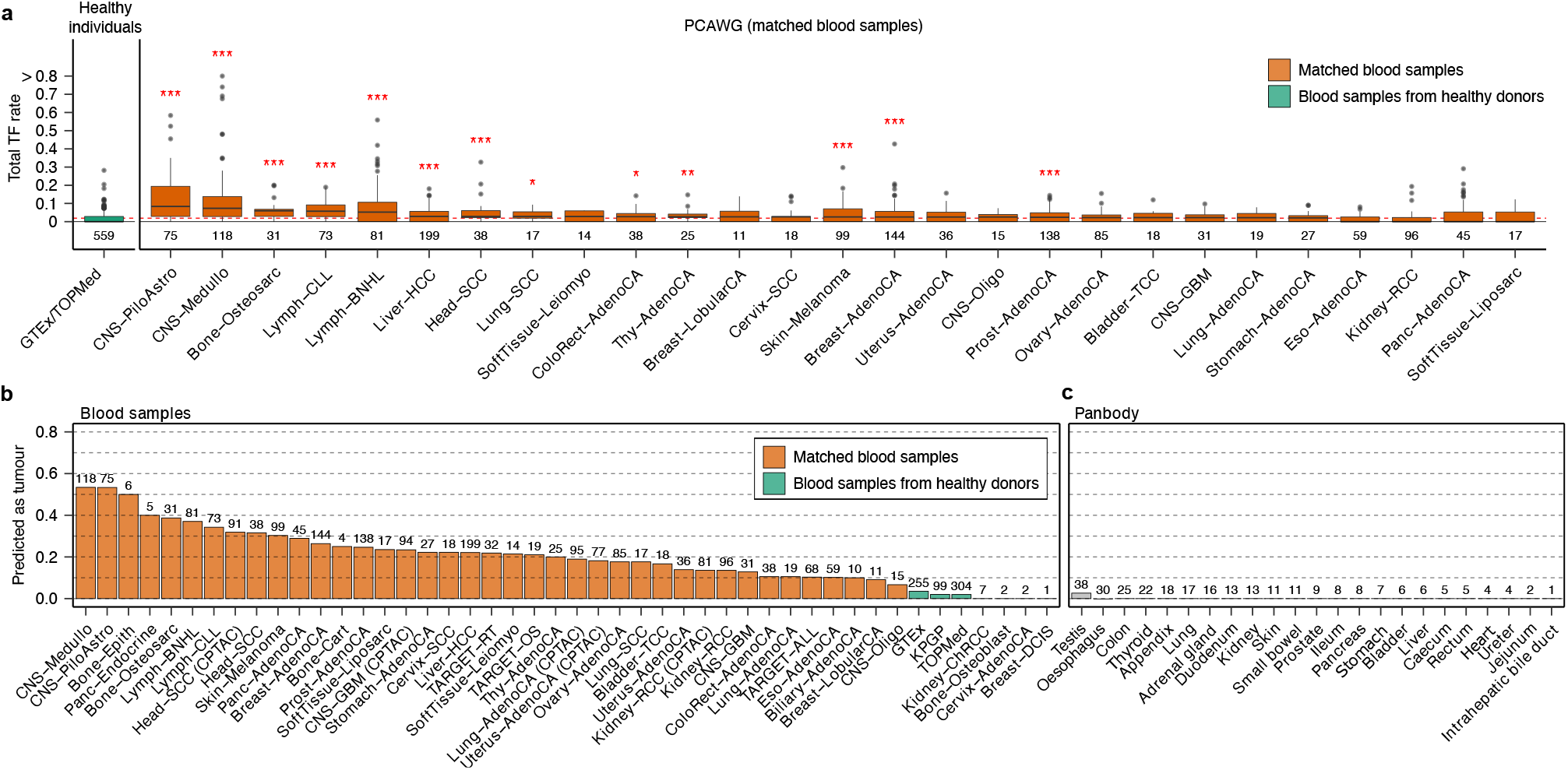
ALT-TFs detected in blood samples enable cancer detection. **a**, ALT-TF rates in blood samples from healthy individuals from GTEx and TOPMed (green) and blood samples from the PCAWG cohort (orange). **b**, Fraction of individuals predicted to have cancer based on the patterns of ALT-TFs detected in blood. Predictions were computed using 100 Random Forest models trained on features of the ALT-TFs detected in WGS data from matched blood samples from CPTAC, PCAWG and TARGET. WGS data from blood samples from GTEx, KPGP and TOPMed were used as controls for model training. Only samples with at least 1 ALT-TF in blood were used for training. **c**, Fraction of histologically normal samples from the Panbody study predicted as cancer using the same model. The numbers below the boxplots in **a**, and above the bars in **b** and **c** indicate the total number of samples in each group. Box plots in **a** show the median, first and third quartiles (boxes), and the whiskers encompass observations within a distance of 1.5x the interquartile range from the first and third quartiles. \*\*\**P* < 0.001; \*\**P* <0.01; \**P* <0.05, Wilcoxon rank-sum test.

Next, we utilized Random Forest (RF) classification to model the probability that an individual has cancer based on the patterns of ALT-TFs detected in blood (Methods). For this analysis, we also included blood samples from 438 cancer patients from the Clinical Proteomic Tumor Analysis Consortium (CPTAC) cohort, 119 pediatric cancer patients from The Therapeutically Applicable Research to Generate Effective Treatments (TARGET) program, and 99 healthy individuals from Korean Personal Genome Project (KPGP)^39^. In addition, we analysed 279 non-neoplastic samples spanning 29 histological units from the Panbody study^40^. In brief, each blood sample, from either a healthy donor or a cancer patient, was encoded by a vector recording 117 features of the ALT-TFs detected (Supplementary Table 6 and Methods). The final RF predictions revealed high sensitivity for medulloblastomas (53%), pilocytic astrocytomas (53%), bone neoplasm epithelioid (50%) and pancreatic adenocarcinomas (40%) (Fig. 4b). These values were notably higher when only considering blood samples with at least 1 ALT-TF and remained comparable across cancer stages (Supplementary Fig. 9b-c). In contrast, the false positive rate was low in all control samples (2-4%) and even lower for samples from the Panbody study (0.36%; Fig. 4b)^40^, highlighting the high specificity of this approach (Supplementary Table 7). The most predictive features included the number of pure ALT-TFs, the total number of ALT-TFs, the length of the breakpoint sequence, and the abundance of the TVRs TGAGGG and TTAGGG, which have been previously linked with ALT activity (Supplementary Fig. 10)^19,22^. Notably, we obtained a comparable sensitivity of detection even for non-ALT tumours (Supplementary Fig. 10d). This is consistent with studies reporting the coexistence of telomerase expression and ALT in the same cell populations *in vitro*^41,42^ and in primary tumours^43–45^. Together, these results indicate that the detection of somatic ALT-TFs in blood represents a highly specific biomarker for liquid biopsy analysis.

## Discussion

Here, we report the discovery of a new type of telomere fusions (ALT-TFs), which we link mechanistically with the activity of the ALT pathway. ALT-TFs permit the identification of the telomere maintenance mechanism of tumours in a quantitative manner using sequencing data, thus representing an alternative approach to current methods^22,46^. Using a non-invasive method to determine the telomere maintenance mechanism of tumours could facilitate the introduction of the ALT-status biomarker in clinical practice, which could inform prognosis and treatment selection^47^.

Two features allow to distinguish ALT-TFs from the conventional TFs joining two chromosomes: (i) the presence of outward fusions, and (ii) their localization to DNA fragments rather than chromosomal regions. In addition, we found that microhomology at the fusion point is common in ALT-TFs. Microhomology results from the annealing of the self-complementary TA sequence, which is contained within canonical T<u>TA</u>GGG telomeric repeats. The relevance of the TA sequence in the formation of telomere fusions is consistent with previous studies indicating an increment of telomere fusions after artificially increasing the self-complementarity of telomere sequences^14^. Since telomere fusions can halt cell division, we infer that avoiding excessive self-complementarity in telomeric repeats would favour tumour expansion. Supporting this view, human tumours contain numerous TVRs that lack self-complementarity. Specifically, the most conspicuous TVRs found in human tumours (T<u>**G**A</u>GGG; T<u>**C**A</u>GGG; T<u>T**G**</u>GGG; T<u>T**C**</u>GGG)^19,30^ destroy the self-complementary TA sequence. In addition, single TVRs are interspersed throughout canonical TTAGGG sequences in cells positive for ALT markers limiting the chances of self-complementary alignments^19^. Based on these observations, we hypothesize that telomere sequences that lack self-complementarity are strongly selected for in tumours in which telomere breakage (deprotection) is favoured, as is the case in ALT positive cells.

We show that ALT-TFs are detected in the blood of cancer patients, which can be used for liquid biopsy analysis with high sensitivity and high specificity. We obtain high sensitivity for tumour types for which early detection methods are severely lacking, such as pancreatic cancer, and tumours that are difficult to sample, such as pediatric brain tumours and glioblastomas. Notably, we report high sensitivity and specificity for predictive models trained on blood WGS data generated using standard protocols for germline genomic analysis. Enrichment or amplification of DNA fragments containing ALT-TFs followed by sequencing or other readouts^48^ could further improve sensitivity and scalability, thus complementing current liquid biopsy methods based on the detection of methylation^49^, point mutations^50^ or genomic instability patterns^51^ in circulating tumour DNA. Overall, detection of ALT-TFs in blood represents a new method for liquid biopsy analysis, with implications for early detection, patient stratification and disease monitoring.

## Methods

### Data sets

To characterize the landscape of somatic TFs in human cancers, we analysed WGS data from 2071 tumour-normal sample pairs from the PCAWG consortium^52^, 306 cancer cell lines from the CCLE^23^, 119 blood samples from the TARGET project, and 438 blood samples from the CPTAC consortium^53,54^. In addition, to assess the rate of TFs in non-cancer samples, we curated WGS data sets from 255 blood samples from the GTEx project^55^, 304 blood control samples from control individuals from the Genetic Epidemiology of COPD project (COPDGene), which is part of the TOPMed Program, 2490 EBV-transformed cell lines from the 1000 Genomes Project^56^, 99 blood samples from the KPGP project^39^, and 279 non-neoplastic samples from the Panbody study (from individuals PD43851/PD42565, PD43850 and PD28690)^40^. Finally, to elucidate the underlying mechanisms of TF formation, we analysed WGS data from 117 retinal pigment epithelium (RPE-1) clones sequenced upon induction of telomere crisis^10^ and 7 controls^9^. The list of data sets, sample IDs, and alignment details for each sample are provided in Supplementary Table 2.

### Detection of telomere fusions using short-read sequencing data

We developed TelFusDetector to detect TFs using sequencing data (https://github.com/cortes-ciriano-lab/telomere_fusions). To identify regions containing fusion-like patterns in the human genome that could be misclassified as somatic, we first scanned the human reference genome (builds hg19, GRCh38 and T2T-CHM13) for patterns of TFs in windows of 500bp with an overlap of 250bp. As expected, this analysis yielded the relic of a telomere fusion in chromosome 2^21^. In addition, we discovered additional regions with forward and reverse telomere repeats in chromosome 9, which we term “chr9 endogenous fusion”. The coordinates for the genomic regions in the reference genome builds hg19, GRCh38 and T2T-CHM13 containing telomeric repeats are provided in Supplementary Table 1.

To detect somatic TFs in sequencing data sets, we extracted aligned sequencing reads containing at least two consecutive TTAGGG and two consecutive CCCTAA telomere sequences using custom python scripts relying on the Pysam and regex modules (https://github.com/pysam-developers/pysam). We allowed for up to two mismatches in the telomeric repeats to consider TVRs^20^. Next, we extracted the mate for each read containing telomeric repeats, and filtered out duplicate reads, as well as supplementary and complementary alignments. Reads mapping to regions in the reference genome containing telomere fusion-like patterns were discounted as somatic.

For the detection of TFs in ALaP-Seq data we used an alternative pipeline that applied FastQC to control for the quality of reads, Cutadapt to remove adapter sequences, and in-house Perl scripts. First, we identified candidate reads containing putative fusions by searching for the presence of two consecutive circularly permuted TTAGGG motifs and two consecutive circularly permuted CCCTAA motifs. Reads mapping to endogenous fusions identified in the mouse reference genome (mm10) were discarded. Finally, we classified the ALT-TFs by analysing the relative orientation of the circularly permuted TTAGGG and CCCTAA repeats and the length of the breakpoint sequence.

### Quality control of telomere fusion calls

To increase the specificity of the ALT-TF calls, we applied a set of filters to remove low-quality read pairs based on alignment quality information and the sequence context of the telomeric repeats. Specifically, we filtered out read pairs with candidate ALT-TFs if (i) at least one read in the pair contained more than one fusion breakpoint of the outward or inward type; (ii) the mate read length was greater than 60bp, did not contain telomeric repeats, and mapped with MAPQ >= 8 to a non-subtelomeric chromosomal region^19^; or (iii) at least one read in the pair mapped to an endogenous TF. The coordinates for telomere regions were downloaded from http://genome.ucsc.edu. Subtelomeric regions were defined as 100 Kbp regions from each chromosomal end. Read pairs supporting the same ALT-TF in each read, as defined by identity of the breakpoint sequence and type of fusion, were considered to originate from fragments with a short insert size and were thus collapsed into a single read. Those read pairs where both mates supported ALT-TFs in the same orientation but with distinct, non-reverse complement breakpoint sequences were discarded.

After applying these filtering criteria, we removed low-quality samples. Specifically, we discarded samples with less than 5 reads mapping to the chromosome 9 endogenous fusion, or if 90% of the reads mapping to this region did not show the expected breakpoint sequence TTAA, which corresponds to the 1% quantile estimated utilizing the PCAWG cohort. In addition, we removed samples with an unusually high rate of filtered reads. Specifically, we discarded the top 1% samples with the highest fraction of sequencing reads with candidate ALT-TFs filtered out in the previous steps.

### Classification of telomere fusions

We classified ALT-TFs into 3 categories based on the relative position of the telomeric sequences TTAGGG and CCCTAA. We classified ALT-TF characterized by 5’-TTAGGG-3’ telomeric repeats followed by 5’-CCCTAA-3’ as inward, and those with 5’-CCCTAA-3’ repeats followed by 5’-TTAGGG-3’ as outward. Finally, read pairs where a read in the pair supports an inward ALT-TF and the other an outward ALT-TF were classified as circular (in-out) events, as this pattern would be expected for ALT-TF formed through ligation and subsequent circularization of short terminal telomere fragments. In addition, we classified ALT-TFs into pure or alternative based on the length and type of sequence at the breakpoint junction. Specifically, we extracted the sequence flanked by the telomeric repeats TTAGGG and CCCTAA allowing for one mismatch or indel in each of the repeats to account for telomere variant repeats and sequencing errors. ALT-TFs with breakpoint sequences in the set of all possible permutations of TTAGGG and CCCTAA were classified as pure, and those with breakpoint sequences longer than 12bp as alternative.

### Estimation of the fragment size of DNA fragments containing ALT-TFs

To assess whether ALT-TFs originate from short DNA fragments, we compared the insert size distribution for read pairs containing ALT-TFs and read pairs mapping outside telomeric regions. For this analysis, we focused on ALaP-Seq paired-end libraries aligned against the mm10 assembly using bowtie2^57^. We estimated the size of the DNA fragments containing ALT-TFs in ALaP-Seq paired-end libraries by assembling reads containing ALT-TFs and their mates using the EMBOSS Merger^58^. Given the low complexity of telomere repeat-containing sequences, we only considered read pairs satisfying the following criteria: (i) the same ALT-TF was detected in both reads in the pair; and (ii) the overlapping region in the assembled sequence was at least 10bp long and with no mismatches, and at least 10% divergent from canonical telomeric repeats, including TVRs. As a baseline for comparison, we estimated the fragment length for read pairs in the same library not mapping to telomeric regions. Alignments were filtered to keep only paired and properly mapped reads. We only considered for further analysis read pairs with an estimated insert size shorter than 190bp given that we require an overlap of at least 10bp in the assembly of read pairs containing ATL-TFs.

### Identification of breakpoint sequences in TFs

To estimate the error rate in reads containing fusions, which limit their accurate identification, we analysed sequencing reads mapping to the chromosome 9 endogenous region in tumours and matched blood samples from PCAWG, as well as in blood samples from GTEx and TOPMed (Supplementary Fig. 1a-d). Because the chromosome 9 endogenous fusion is fixed in the population, only one breakpoint sequence should be detected in each sample (i.e., TTAA), and any additional breakpoint sequences would be the consequence of sequencing errors in the flanking TTAGGG or CCCTAA repeats. Using this approach, we estimated a base error rate at telomeric repeats of 0.74%, which is comparable to the expected sequencing error rate for Illumina data^59^. We detected at least two distinct breakpoint sequences (including TTAA) in 1725 of the 2163 samples (79.8%), indicating the relevance of accounting for sequencing errors for the accurate characterization of ALT-TF breakpoints. Using the reads mapping to the chr9 endogenous fusion, we developed a set of rules for error correction to account for mismatches and indels in the flanking telomeric repeats available at ttps://github.com/cortes-ciriano-lab/telomere_fusions. This correction step reduced the average number of distinct breakpoint sequences for the chr9 endogenous fusion to 1.07 and identified the breakpoint sequence TTAA only in 93.3% of cases (Supplementary Fig. 1b).

### Calculation of telomere fusion rates

To compare the number of ALT-TFs across samples aligned against difference builds of the human reference genome and sequenced using varying sequencing read length and depth, we computed a telomere ALT-TF rate for each sample as:

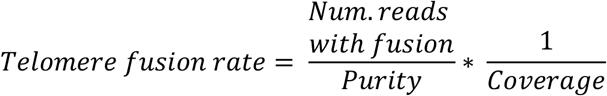

where

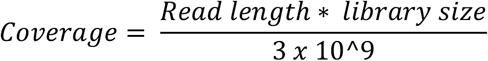

and *Library size* corresponds to the number of mapped reads in a sample after excluding duplicate reads and supplementary alignments. The purity values were obtained from PCAWG marker study^52^. To calculate the rate of endogenous TFs in matched normal samples and cancer cell lines we used a purity value of 1.

### Detection of telomere fusions using long-read sequencing

Due to the higher error rate of long-read sequencing as compared to Illumina sequencing, we required at least 15 TTAGGG and 15 CCCTAA telomeric repeats and at least 5 TTAGGGTTAGGG and 5 CCCTAACCTAA repeats to make a telomere fusion call. Sequencing reads mapping to non-subtelomeric regions with a mapping quality >= 8 and supplementary alignments were discarded. Fusion counts were then normalised for each sample by dividing the number of reads with ALT-TFs by the total number of reads.

### Identification of genomic correlates of ALT-TF formation

To investigate the genomic correlates of ALT-TF formation, we first used a linear model to regress out the effect of the cancer type and tumour purity on the rate of ALT-TFs. We limited this analysis to tumour samples from the PCAWG project with a tumour purity of at least 30%. The residuals of the linear model were then used in a second linear model that incorporated age at diagnosis and sex as covariates, as well as other variables related to genome instability and telomere maintenance. Specifically, we encoded using categorical variables the presence of whole-genome doubling events^52^ and chromothripsis in at least one chromosome^60^. Tumour ploidy and purity were encoded as continuous variables. We also included as categorical covariates in the model the presence of activating mutations or SVs in *TERT* and the ALT pathway classification reported by de Nonneville et al.^22^ or Sieverling et al.^19^. The telomere content normalized against the matched normal sample was included as a continuous variable^19^. Finally, due to the relationship between EBV transformation and ALT activity, we also included the presence of viral sequence insertion events in the tumour genome as a categorical variable^61^. We performed backward stepwise model selection using the Akaike information criterion to identify covariates associated with ALT-TF formation.

### Machine learning

We utilized the R package *RandomForest* to generate Random Forest (RF) models. We used default parameter values for model training, except for the number of trees, which was set to 100. To assess the robustness of the predictions, we trained a total of 100 RF models, each time assigning a different set of samples to the training and test sets. We selected as the optimal cut-off value the probability that maximized sensitivity and specificity across the 100 models. The predictive power of each covariate was estimated by computing the change in accuracy when excluding each covariate from the training data using the function *importance* from the *RandomForest* package.

### ALT prediction using ALT-TFs detected in tumour samples

We generated RF models for the prediction of the ALT status of tumours using the patterns of ALT-TFs detected in cancer samples as covariates. As a training set, we considered as ALT those tumours positive for the C-circle assay, or classified as ALT by both de Nonneville et al.^22^ and Sieverling et al.^19^ We then randomly assigned two-thirds of the ALT-positive samples and an equal number of non-ALT samples from cancer patients to the training set, and held out the rest for testing the model. We encoded the ALT-TFs detected in each sample using a set of 117 variables, which are listed in Supplementary Table 6.

### Cancer detection using ALT-TFs identified in blood samples

We trained RF models for the detection of cancer using the patterns of ALT-TFs detected in blood samples. For model training we focused on samples with at least 1 ALT-TF; individuals with no ALT-TFs detected in blood were considered to be cancer-free. We randomly assigned two-thirds of the GTEx, TOPMed and KPGP samples and an equal number of blood samples from cancer patients from the PCAWG, CPTAC and TARGET projects to the training set, and held out the remaining 1296 samples for testing the model. We encoded the ALT-TFs detected in each sample using the same set of 117 variables used to predict the ALT status of tumours (Supplementary Table 6). These include the fraction of each TVR in the reads supporting ALT-TFs and ALT-TF rates. Finally, the predictions of cancer status were obtained by selecting the most frequent prediction across 100 models, each trained on a random subset of the training data. As additional validation, we applied the obtained RF models to a set of 29 diverse histological structures from normal tissue samples from the Panbody study to investigate the specificity of our approach.

## Data availability

The raw sequencing data from the PCAWG project is available through controlled access application to the International Cancer Genome Consortium Data Access Compliance Office (DACO; http://icgc.org/daco) for the ICGC portion, and to the TCGA Data Access Committee (DAC) via dbGaP for the TCGA portion (https://dbgap.ncbi.nlm.nih.gov/aa/wga.cgi?page=login; dbGaP Study Accession: phs000178.v1.p1). Additional information on accessing the PCAWG data, including processed data sets, can be found at https://docs.icgc.org/pcawg/data/. We used the following data sets from PCAWG, which are available at Synapse (https://www.synapse.org/) under the Synapse IDs syn10389158: clinical data from each patient, including tumour stage and vital status; syn1038916: harmonised tumour histopathology annotations using a standardised hierarchical ontology; and syn8272483: purity and ploidy values for each tumour sample. The raw sequencing data generated by the GTEx project is available through controlled access application via dbGAP (dbGaP Study Accession: phs000424.v8.p2). The raw sequencing data generated by the Genetic Epidemiology of COPD (COPDGene) project, which is part of TOPMed, is available through controlled access application via dbGAP (dbGaP Study Accession: phs000951.v4.p4). The raw sequencing data from the Korean Personal Genome Project (KPGP) were downloaded from SRA (study accession: PRJNA284338). Raw sequencing data from the CPTAC3 study is available through controlled access application to the NCI Data Access Committee (DAC) via dbGAP (dbGaP Study Accession: phs001287.v5.p4). WGS data from 124 retinal pigment epithelium (RPE) clones were downloaded from the European Nucleotide Archive database under primary accession number PRJEB23723^10^ and European Genome-Phenome Archive (EGA, hosted by the EBI and the CRG) under the accession number EGAD00001001629^9^. The raw sequencing data from the Panbody project were downloaded from EGA (EGAD00001006641). PacBio and Illumina WGS data from the breast cancer cell line SK-BR-3 were downloaded from http://schatz-lab.org/publications/SKBR3/^24^. Nanopore and Illumina WGS data from the melanoma COLO829 cell line were downloaded from the European Nucleotide Archive (ENA) under the study accession PRJEB27698^25^ PacBio WGS data from the colon cancer cell lines HCT116, KM12, SW620 and SW837 are available at the Gene Expression Omnibus (GEO) database under the accession number GSE149709^26^. AlaP-Seq and CHIRT-seq data are available at GEO under accession numbers GSE135563 and GSE79180, respectively.

## Code availability

The code to detect ATL-TFs using sequencing data is available at: https://github.com/cortes-ciriano-lab/telomere_fusions

## Author contributions

I.C.-C. and I.F designed and supervised the study. F.M. and M.J.G.R. performed analyses. F.M. generated the figures with input from M.J.G.R., I.C.-C., and I.F. F.M. and I.C.-C. implemented TelFusDetector with input from M.J.G.R. and I.F. F.M., I.C.-C., and I.F. wrote the manuscript with input from M.J.G.R. All authors read and approved the final manuscript.

## Acknowledgements

F.M. and I.C.-C. thank EMBL for funding. I.F. was funded by grants from the Spanish Ministry of Science and Innovation (PID2019-110339RB-I00) and the Comunidad de Madrid (S2017/BMD-3875). The CNIC is supported by the Ministerio de Ciencia, Innovación y Universidades and the Pro CNIC Foundation, and is a Severo Ochoa Center of Excellence (SEV-2015-0505). All authors thank the computational resources provided by the European Bioinformatics Institute (EMBL-EBI).

## Supplementary Figures

**Supplementary Figure 1.**
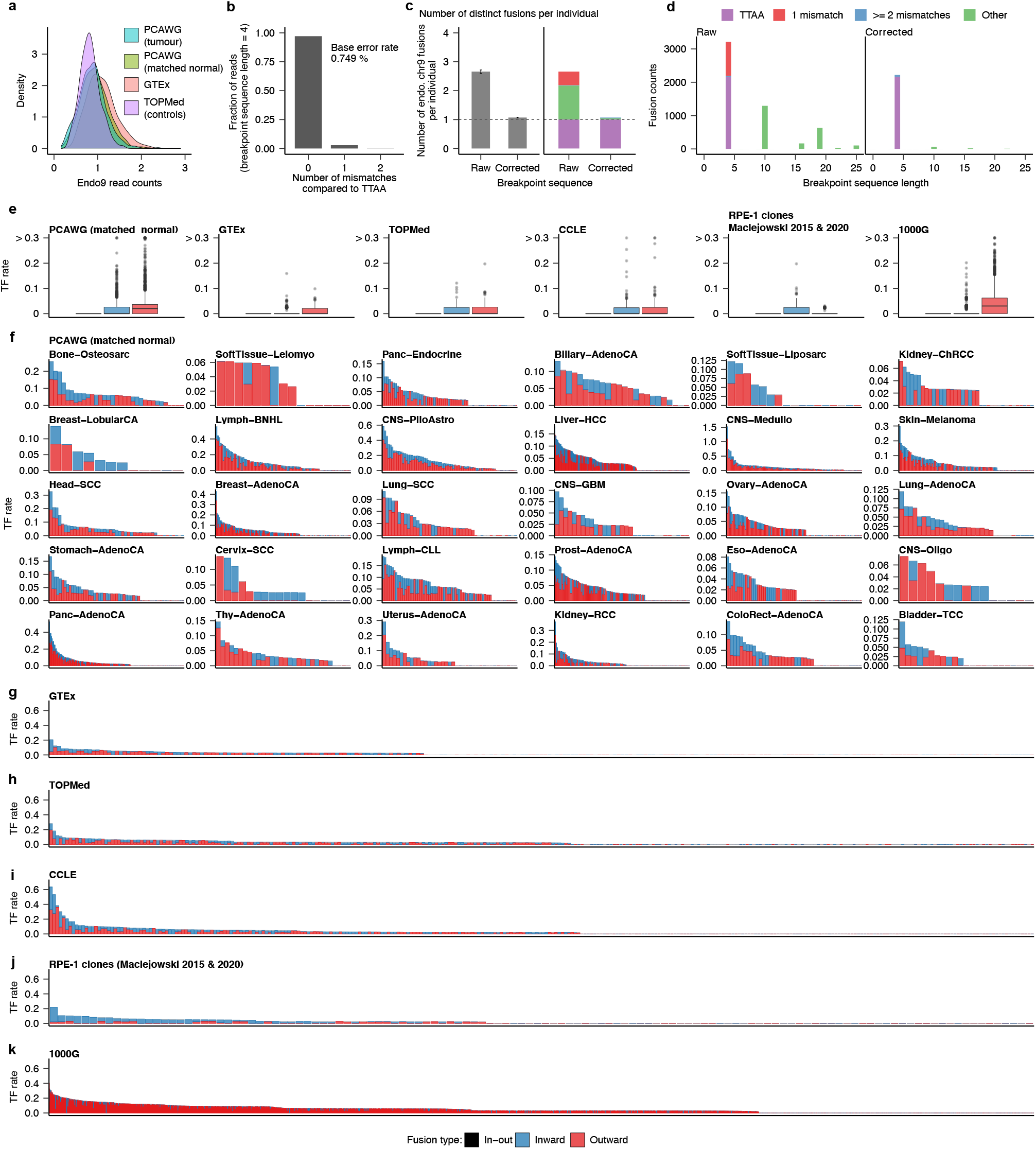
**a**, Distribution of the rate of endogenous chromosome 9 fusions detected in PCAWG tumour and matched-normal samples, and healthy blood samples from GTEx and TOPMed. Overall, the distribution is comparable across projects, indicating that TF rates enable analysis across samples sequenced using different read lengths and at variable sequencing depth. **b**, Fraction of reads mapping to the endogenous chromosome 9 fusion. The bars indicate the fraction of reads with the expected breakpoint sequence TTAA (shown in purple), one mismatch (red), and two mismatches (blue). Only reads with a fusion breakpoint sequence of 4bp were included in this plot. This analysis revealed an error rate in reads mapping to the chromosome 9 endogenous fusion of 0.749%, which is comparable to the expected error rate for Illumina sequencing. **c**, Number of distinct chromosome 9 endogenous fusions (as determined by the breakpoint sequence) found per individual before and after error correcting the breakpoint sequences (Methods). As expected, we detected one breakpoint sequence at the chromosome 9 endogenous fusion after error correction in most samples. **d**, Length of the breakpoint sequences for the endogenous chromosome 9 fusion detected per individual before and after error correction. As expected, only one breakpoint sequence of 4bp in length (TTAA) is detected in most samples. **e**, Distribution of circular, inward and outward ALT-TFs across all samples. ALT-TF rates detected in matched-normal samples from PCAWG (**f**), GTEx (**g**), TOPMed (**h**), cancer cell lines (CCLE) (**i**), RPE-1 cells (**j**), and EBV-immortalized B cell lines from the 1000 Genome Project (**k**).

**Supplementary Figure 2.**
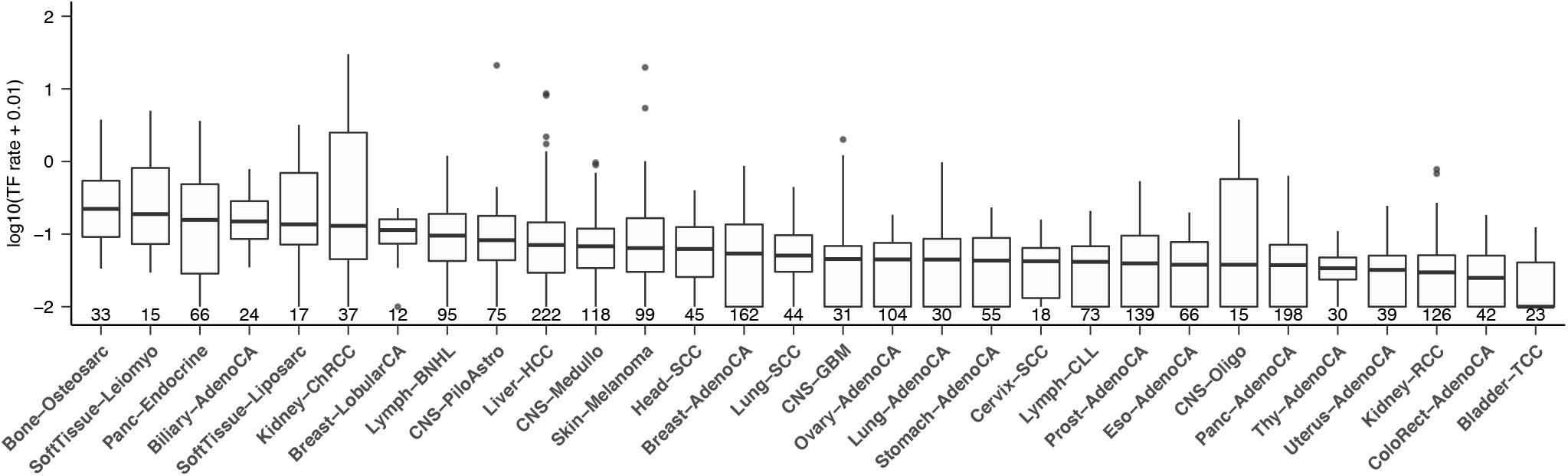
**a**, TF rates across cancer types in the PCAWG cohort. Box plots show the median, first and third quartiles (boxes), and the whiskers encompass observations within a distance of 1.5x the interquartile range from the first and third quartiles.

**Supplementary Figure 3.**
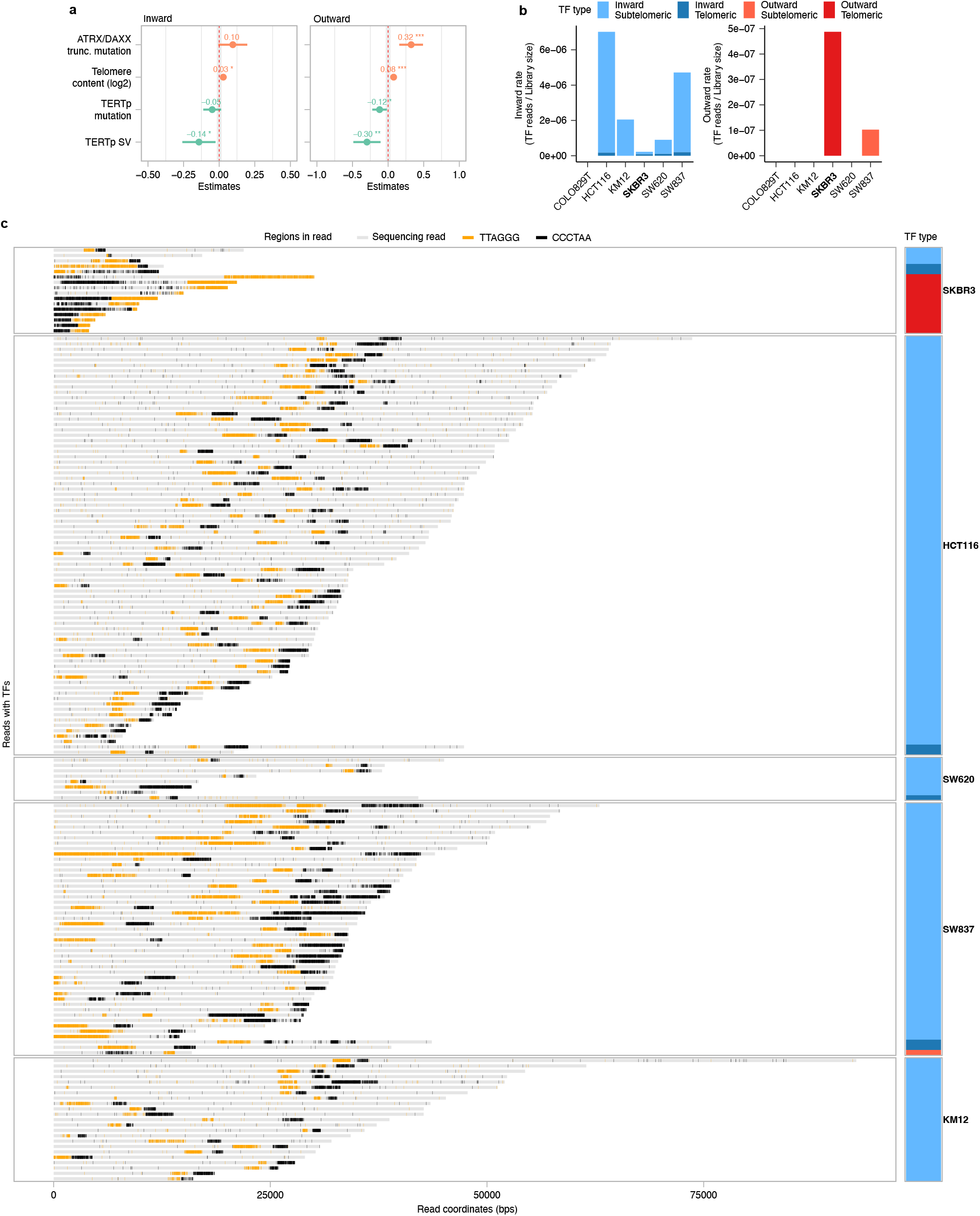
**a**, Coefficient values estimated using linear regression analysis and variable selection for the covariates with the strongest positive and negative association with TF rates. For this analysis we used the ALT status classification reported by Sieverling et al. 2020 (see also Fig. 2a). **b**, TF rates normalised by the total library size obtained for cell lines analysed using long-read sequencing technologies. The rate of outward ALT-TFs is significantly higher in the breast cancer cell line SKBR3, which is a used model of ALT. ALT-TFs were classified as subtelomeric if the genomic regions flanking the telomeric repeats could be unambiguously mapped to a subtelomeric region. **c**, Schematic representation of the long reads containing ALT-TF fusions. The bar on the right represents the classification of the different fusions as described in (**b**). Intra-read telomeric repeats TTAGGG and CCCTAA are shown in orange and black, respectively.

**Supplementary Figure 4.**
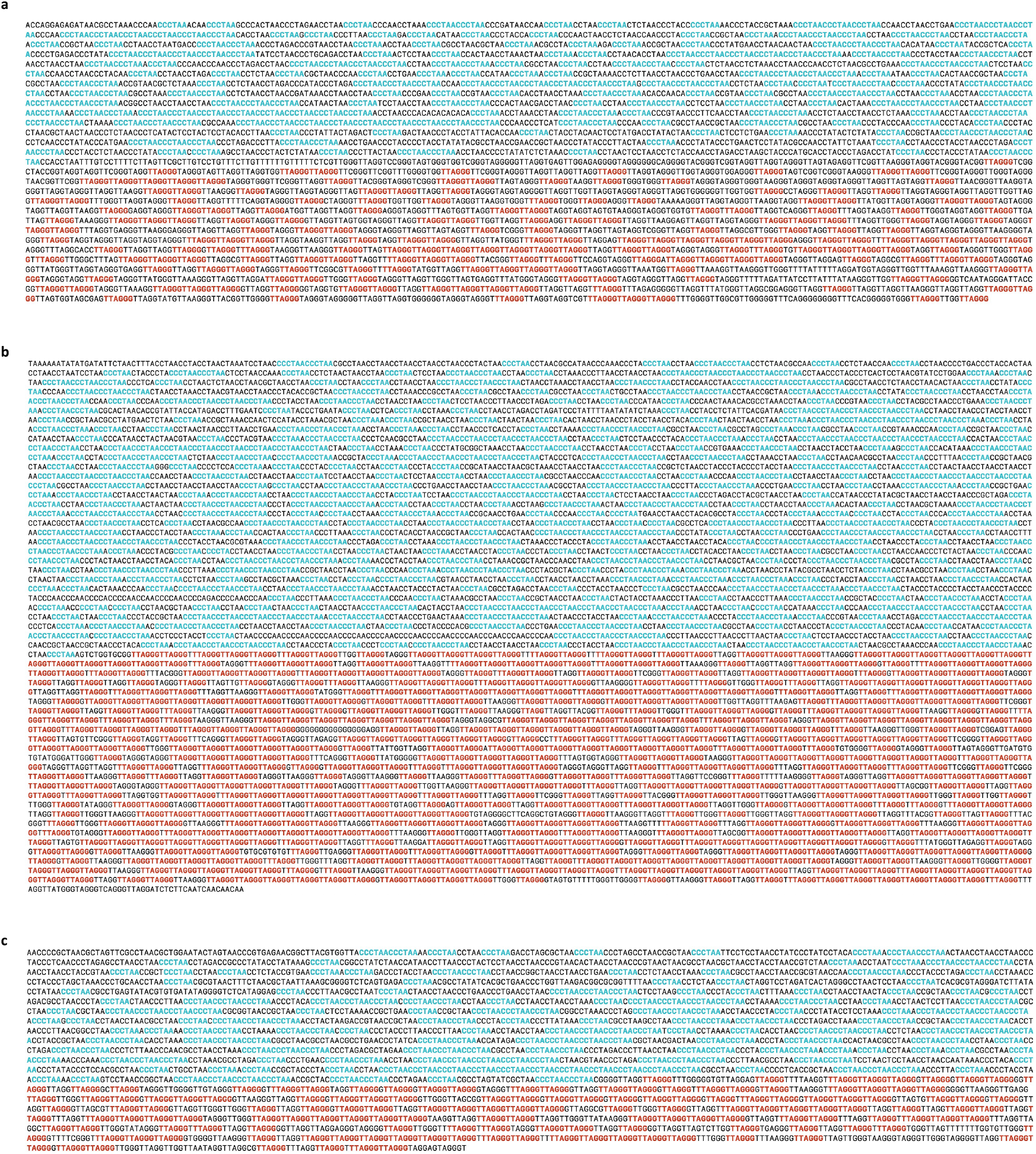
**a-c**, Representative examples of outward fusions detected in PacBio long-read sequencing data from the breast cancer cell line SKBR3. Telomeric repeats TTAGGG and CCCTAA are shown in red and blue, respectively.

**Supplementary Figure 5.**
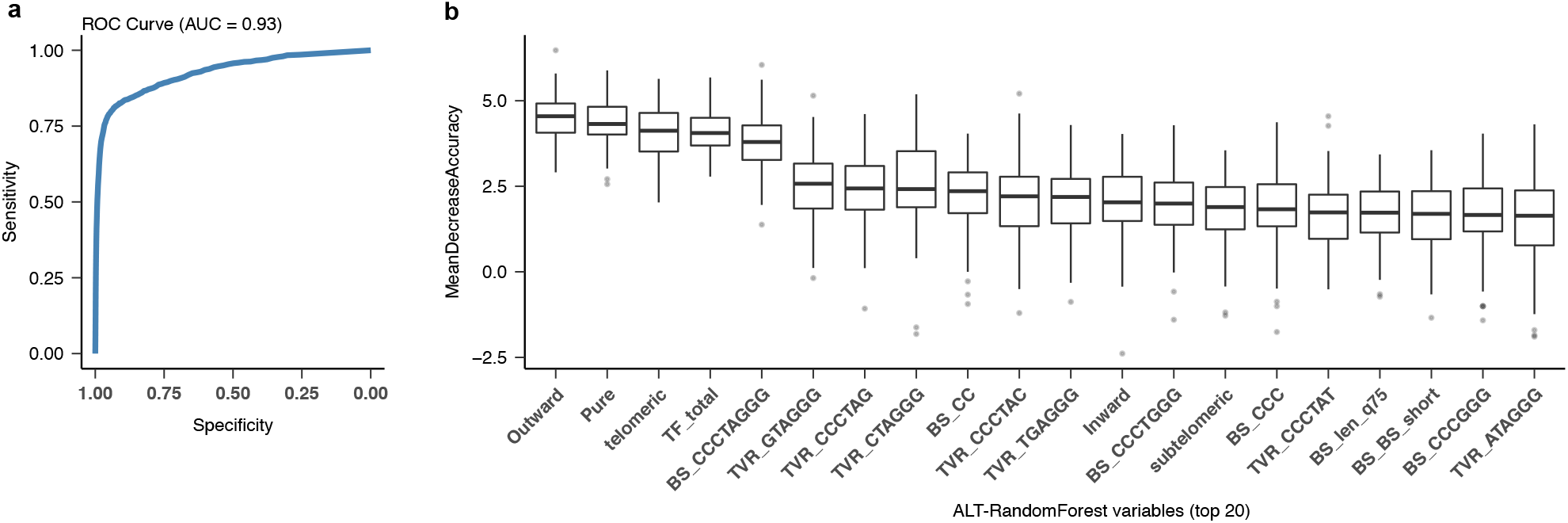
Predictive power of the Random Forest models trained to predict the ALT status of tumours based on the features of ALT-TFs. **a**, Area Under the Curve (AUC) for the Random Forest model. The results across 100 bootstrap resamples are shown. **b**, Mean decrease in accuracy obtained for the variables used in the Random Forest model. Only the top 20 predictive features are shown in the plot. The higher the value the stronger the predictive power of each variable. Box plots show the median, first and third quartiles (boxes), and the whiskers encompass observations within a distance of 1.5x the interquartile range from the first and third quartiles.

**Supplementary Figure 6.**
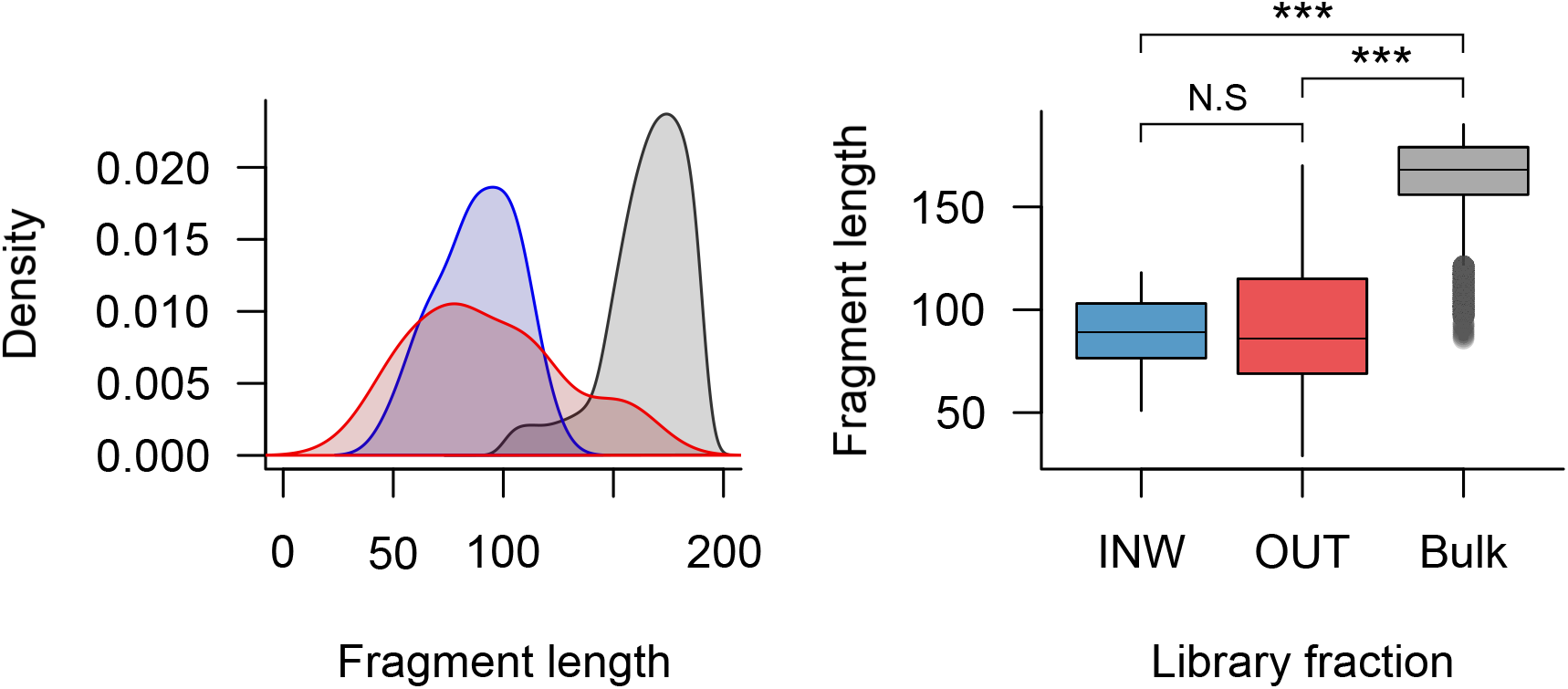
Fragment size distribution for assembled fragments in ALaP-Seq libraries from mESCs with peroxidase corresponding to read pairs mapping outside telomeric regions and read pairs containing either inward (INW, blue) or outward (OUT, red) fusions. The distribution of assembled fragment lengths in complete libraries (Bulk, grey) has a maximum of around 180 bp, while those for inward and outward ALT-TF-containing fractions have maximum sizes below 100 bp (\*\*\**P* < 0.001; Wilcoxon rank-sum test).

**Supplementary Figure 7.**
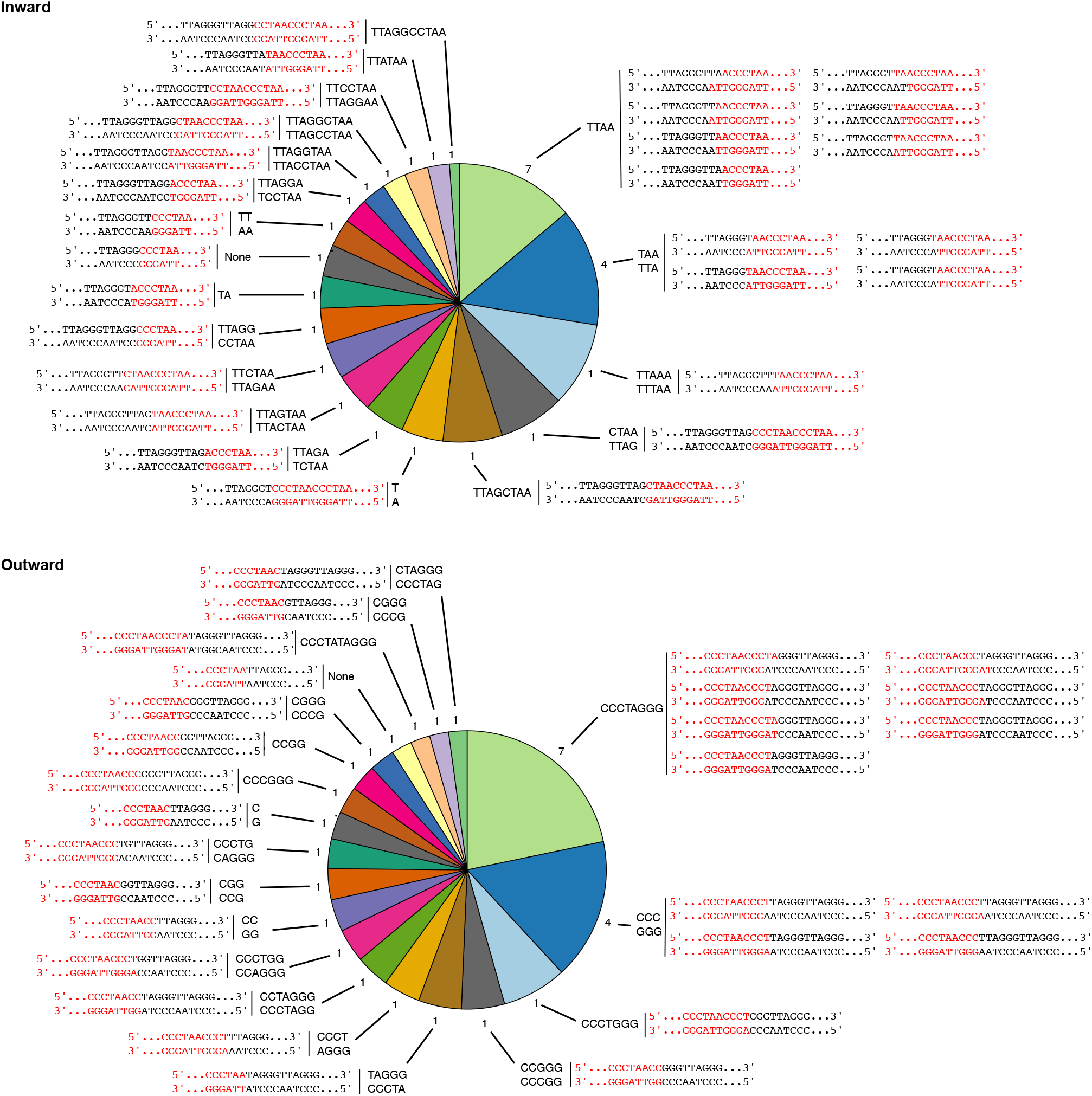
Extended pie chart showing the distribution of the distinct breakpoint sequences observed in pure ALT-TFs in PCAWG tumours. The numbers around the pie charts represent the number of combinations of circular permutations of telomeric repeat motifs that can generate each breakpoint sequence. The legend reports the breakpoint sequences in both strands unless they are identical (e.g., TTAA).

**Supplementary Figure 8.**
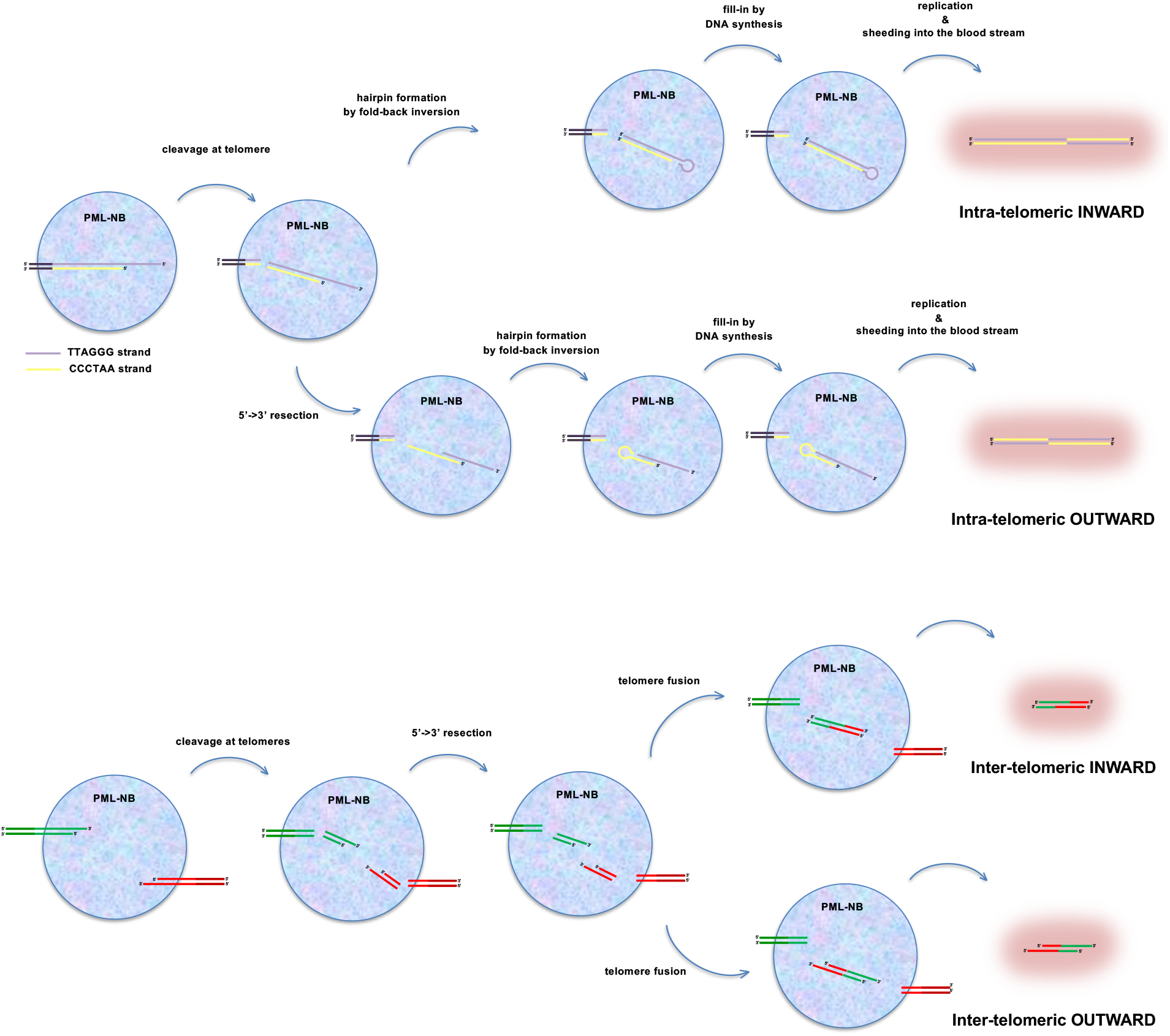
Intra- and inter-telomeric ligation mechanisms in PML-NBs can lead to the formation of ALT-TF detected in the blood of cancer patients. See main text for details.

**Supplementary Figure 9.**
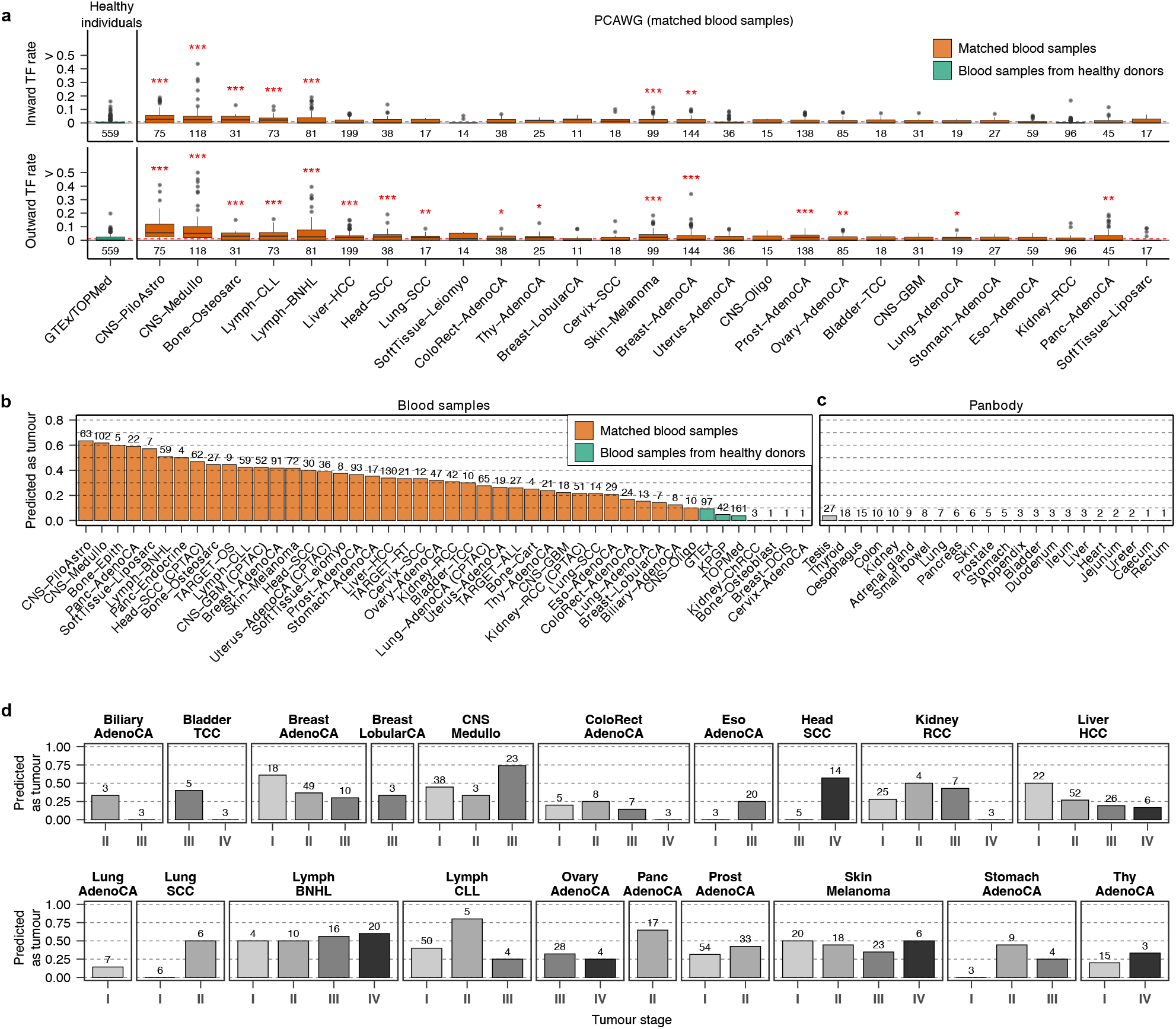
**a**, Rates of inward and outward ALT-TFs detected in the blood samples from healthy individuals from GTEx and TOPMed (green) and matched blood samples from PCAWG (orange). **b**, Fraction of individuals with at least 1 ALT-TF in blood predicted to have cancer. **c**, Fraction of histologically normal samples from the Panbody study with at least 1 ALT-TF predicted as cancer samples using the same model. **D**, Fraction of PCAWG cases with at least 1 ALT-TF in blood correctly classified as having cancer stratified according to cancer stage. Predictions in **b**-**d** were computed using 100 Random Forest models trained on features of the ALT-TFs detected in the WGS data from the matched blood samples from PCAWG, CTPAC and TARGET, and blood samples from GTEx, KPGP and TOPMed, which were used as controls. Only samples with at least 1 ALT-TF in blood were used for training. The numbers below the boxplots in **a**, and above the bars in **b**-**c** indicate the total number of samples in each group. Box plots in **a** show the median, first and third quartiles (boxes), and the whiskers encompass observations within a distance of 1.5x the interquartile range from the first and third quartiles. \*\*\**P* < 0.001; \*\**P* <0.01; \**P* <0.05, Wilcoxon rank-sum test.

**Supplementary Figure 10.**
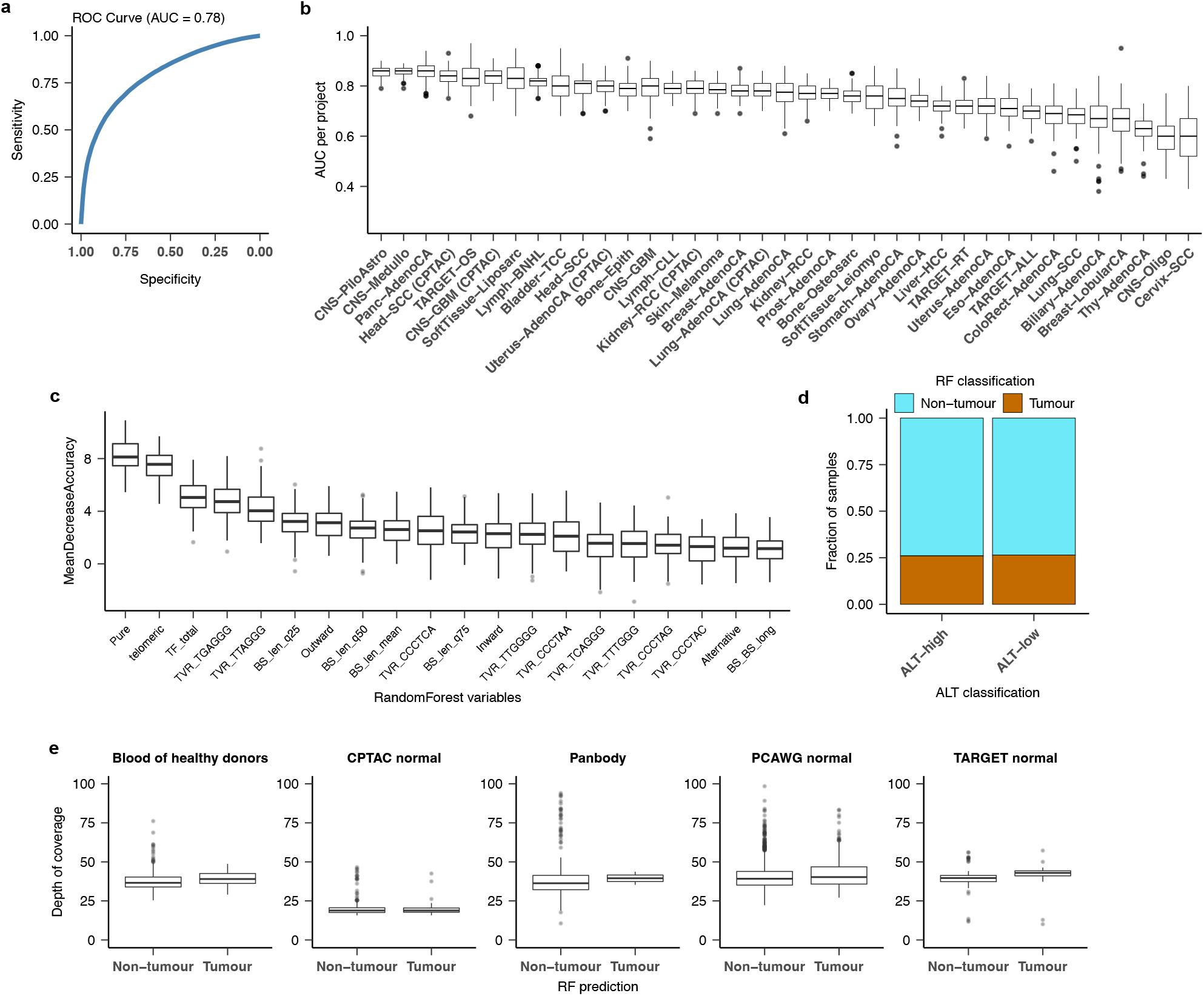
Predictive power of the Random Forest models generated to predict the probability that an individual has cancer on the basis on the features of the ALT-TFs detected in blood. **a**, Global AUC of the Random Forest model. The results across 100 bootstraps are shown. **b**, AUC values split by cancer type. **c**, Mean decrease in accuracy (importance) obtained for each variable used in the Random Forest model. The top 20 predictive features are shown. **d**, Fraction of samples predicted as originating from a cancer patient (“Tumour”) and grouped by ALT classification status. **e**, Depth of coverage of samples predicted to belong to a cancer patient (represented by “Tumour”) or not (“Non-Tumour”). Overall, this analysis indicates that the predictive power of the algorithm is not affected by the sequencing depth. Box plots show the median, first and third quartiles (boxes), and the whiskers encompass observations within a distance of 1.5x the interquartile range from the first and third quartiles.

## Supplementary Tables

**Supplementary Table 1.** The coordinates for the endogenous telomere fusion patterns detected in the human reference genome are listed.

**Supplementary Table 2.** Telomere fusion rates for all samples analysed.

**Supplementary Table 3.** Results of the ANOVA models generated to test the association between ALT-TF rates and the expression of TERRA and TERT.

**Supplementary Table 4.** Analysis of the length of the fragments containing ALT-TFs.

**Supplementary Table 5.** List of breakpoint sequences detected in pure ALT-TFs.

**Supplementary Table 6.** List of 117 variables used to train the Random Forest models designed to predict the ALT status of tumours, and for the detection of cancer based on the features of the ALT-TFs detected in blood samples.

**Supplementary Table 7.** Summary of the Random Forest model results used to predict cancer status.

